# Meditation Styles Are Highly Discriminable from EEG at the Subject Level With Limited Generalization Across the Population: A Machine-Learning Study

**DOI:** 10.64898/2026.05.15.725404

**Authors:** Saqib Hayat, Francesco Goretti, Rachele Fabbri, Chiara Noferini, Elena Cravero, Paolo Mori, Alessandro Scaglione, Francesco S. Pavone

## Abstract

Meditation has been associated with improvements in attention, emotional regulation, and mental well-being, motivating increasing interest in objective methods for assessing meditative states. In this study, we investigate whether EEG-based machine learning can reliably distinguish between multiple meditation styles and mind-wandering states. EEG data were recorded from experienced meditators performing three meditation styles, Shamatha, Vipassana, and Metta, together with an eyes-closed mind-wandering condition. EEG signals were preprocessed to remove artifacts, and features were extracted from frequency, time-frequency, and time domains. Classification was evaluated using both intra-subject and inter-subject strategies with multiple machine learning classifiers. Results demonstrate high intra-subject classification accuracy across meditation-versus-mind-wandering and meditation-style comparisons, indicating strongly discriminative subject-specific neural signatures. In contrast, inter-subject performance decreased substantially, particularly for distinguishing meditation styles, suggesting considerable inter-individual variability in meditation-related EEG patterns. Furthermore, temporal analysis revealed that classification performance increase over time, indicating that the neural distinctions between meditation states become increasingly pronounced over time. Additionally, t-SNE visualization showed clear within-subject clustering but increased overlap across subjects, explaining the reduced inter-subject generalization. Overall, these findings highlight the potential of EEG-based machine learning for personalized assessment and monitoring of meditative states while emphasizing the challenges of developing subject-independent meditation classification systems.

## INTRODUCTION

Meditation is emerging as a set of exercises that helps in the self-regulation of thoughts, emotion, and attention (Young et al., 2021). A number of studies have shown positive effects on mental health of meditation practice, including a general reduction in stress and level of propensity to depression (Chiesa et al., 2011; Saeed et al., 2019). Although meditation encompasses a wide range of practices with distinct cognitive and emotional effects, they are often broadly categorized into different styles based on attentional focus and regulatory mechanisms (Lutz et al., 2008). Moreover, evidence suggests that experienced meditators exhibit greater depth and frequency of sustained meditation, along with reduced frequency and duration of mind-wandering episodes compared to novices (Brandmeyer and Delorme, 2018). Mind-wandering, characterized by spontaneous, task-negative thoughts, emerging when attention is not focused, can interrupt the process of meditation (Sood and Jones, 2013; Hasenkamp et al., 2012). Therefore, it is essential to assess the effectiveness of meditation practice and provide feedback to the meditators to enhance their performance and maximize the mental health benefits (Yu et al., 2022). Consequently, meditation state recognition is of great importance for online or offline feedback during practice to accelerate the learning process and facilitate the attainment of deeper meditative states (Brandmeyer and Delorme, 2013).

Exploring meditation is challenging given the vast heterogeneity in techniques and underlying philosophies (Fox et al., 2016). Nonetheless, the regulation of attention is a central feature of various meditation methods (Davidson, 1976). Meditation practices are commonly categorized into focused attention meditation (FAM) and open monitoring meditation (OMM) based upon direction of attentional process engagement (Lutz et al., 2008). FAM, known as Shamatha in Tibetan Buddhism, involves sustained attention on a chosen object (e.g., breath), while OMM, also known as Vipassana, entails non-reactive awareness of experience without fixation. Both practices are widely used in mindfulness-based interventions and involve the intentional activation, direction, and maintenance of attention, strengthening awareness of thoughts, emotions, and bodily sensations from a neuropsychological perspective (Lutz et al., 2008). Compassion-based Metta meditation, emphasizes emotional cultivation through loving-kindness (Hofmann et al., 2011a). Shamatha and Vipassana are widely employed for attentional regulation and in clinical interventions for depression, anxiety, and stress (Jerath et al., 2015), whereas Metta has shown efficacy in cognitive-behavioral therapy for chronic depression (Hofmann et al., 2011b). These distinct yet complementary styles were selected in our study for their relevance to both cognitive and affective processes, enabling a comprehensive investigation of meditative state effects.

The meditative state has been investigated using various neuroimaging and neurophysiological techniques, including magnetic resonance imaging (MRI) (Gotink et al., 2016), functional magnetic resonance imaging (fMRI) (Engström et al., 2022), functional near-infrared spectroscopy (fNIRs) (Xie et al., 2022), electroencephalogram (EEG) (Neri et al., 2024; Shang et al., 2023; Ahani et al., 2014). Among these, EEG remains the most widely used non-invasive method for capturing meditation-related brain activity due to its high temporal resolution.

Meditation increases overall brain activity (Lee et al., 2018), and the effects of short-term Shamatha (Cahn and Polich, 2006; Neri et al., 2024), Vipassana (Cahn et al., 2010, 2013; Lomas et al., 2015), and Metta (Lutz et al., 2004; Hofmann et al., 2011a) meditation on EEG have been observed. The effects of these meditation practices on EEG are commonly reported in the spectral domain across five frequency bands, namely delta (1–4 Hz), theta (4–8 Hz), alpha (8–12 Hz), beta (13–30 Hz), and gamma (30–45 Hz). The most common effects of meditation are observed in the theta and alpha bands, although findings are not consistently reported across studies (Cahn and Polich, 2006; Lomas et al., 2015). FAM and OMM are associated with increases in anterior theta and posterior alpha activity (Lee et al., 2018). Additionally, occipital gamma activity has been observed in experienced meditators during Vipassana practice (Braboszcz et al., 2017; Cahn et al., 2010), as well as in FA and OM-based Buddhist meditation (Luft et al., 2019). Moreover, different meditation styles exhibit distinct neural effects due to their unique cognitive and attentional characteristics (Lee et al., 2018).

Machine learning approaches have increasingly demonstrated their ability to extract meaningful patterns from high-dimensional and complex biological datasets, including genomics (Libbrecht and Noble, 2015; Iadanza et al., 2022), neuroimaging (Davatzikos, 2019), optical neural recordings (Scaglione et al., 2022) and electrophysiological recordings (Lotte et al., 2018). Advances in computational analysis have thus enabled automated classification of meditative states using machine learning methods and EEG features. EEG features are typically derived from frequency (Banquet, 1973; Neri et al., 2024), time-frequency (Tee et al., 2020), or time-domain (Diykh et al., 2016) analyses. These features are then input into machine learning models—such as LDA (Panachakel et al., 2021a), SVM (Ahani et al., 2014; Shaw and Routray, 2016a), and Random Forest (Huang et al., 2021)—as well as deep learning models, including convolutional neural networks (CNNs) (Shang et al., 2023) and long short-term memory (LSTM) networks (Panachakel et al., 2021b), to reliably distinguish meditation states. Traditional machine learning methods are typically applied to handcrafted features, while deep learning approaches can be applied either to extracted features or directly to raw EEG data, enabling automatic learning of complex representations from large datasets (Craik et al., 2019). In particular, many CNN-based approaches use EEG spectrograms as image inputs, transforming signal representations into learnable visual patterns for classification (Vrbancic and Podgorelec, 2018). Feature extraction-based approaches remain more common for small datasets, whereas raw-data-driven deep learning methods are generally preferred when large-scale datasets are available (Craik et al., 2019).

Past research on meditation state classification using EEG and machine learning has primarily focused on inter-state differences (meditation vs. rest) (Shang et al., 2023; Ahani et al., 2014; Panachakel et al., 2021b), inter-group differences (meditation practitioners vs. novices) (Shaw and Routray, 2016b), meditation experience-based classification (Lee et al., 2017), and the effects of meditation (Jadhav et al., 2016; Gupta et al., 2021). Most of these studies rely on a single type of meditation. There is still a lack of research on meditation state classification involving multiple meditation traditions, particularly the widely practiced three styles, and to the best of our knowledge, no study has investigated brain state classification using Shamatha, Vipassana, and Metta meditation within a unified framework. Unlike conventional binary classification frameworks (e.g., meditation vs. rest), the inclusion of multiple meditation styles enables finer-grained characterization of distinct neural signatures associated with different attentional and affective processes, thereby reducing the risk that classification performance is driven by meditation-specific features that may not generalize across techniques.

In this study, we aim to distinguish between multiple meditative states using machine learning methods to enable reliable assessment of meditation. We hypothesize that each meditation style induces state-specific changes, resulting in EEG features that are significantly different from one another and from those observed during mind-wandering. Furthermore, we hypothesize that these features will enable high discrimination of meditation styles at the subject level and may also generalize across individuals. To test this hypothesis, volunteer subjects performed the above-mentioned three meditation practices, preceded by an eyes-closed mind-wandering baseline. Using these datasets, we conducted analyses focusing on: (1) robust extraction and selection of EEG features that characterize meditative states, (2) comparative evaluation of high-performance machine learning models in intra- and inter-subject settings, and (3) temporal analysis to capture dynamic changes over time.

## MATERIAL AND METHODS

The pipeline of the methodology consists of data recording, data preprocessing, feature extraction, feature selection and meditation state recognition using machine learning algorithms (see Figure 1).

**Figure 1.**
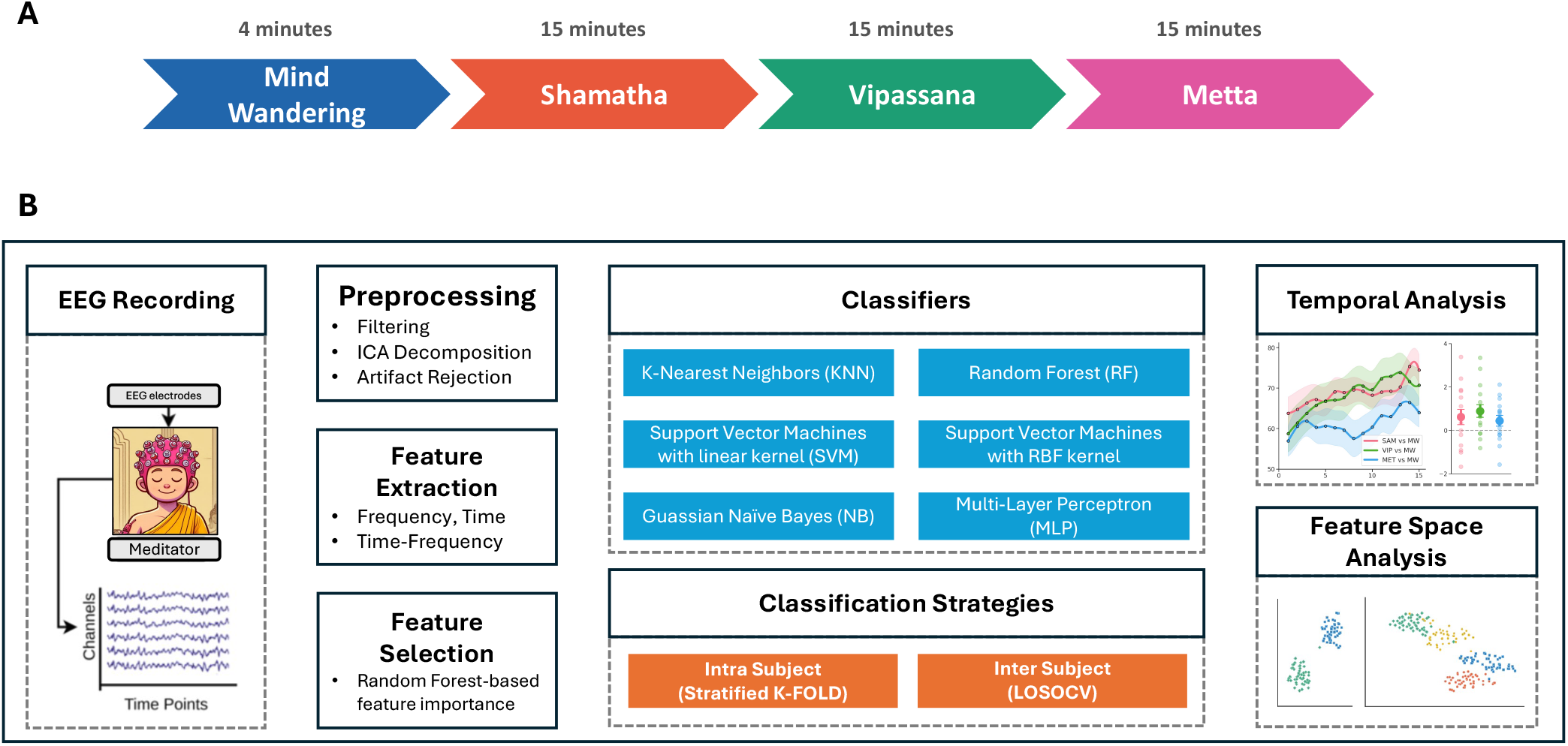
Flow diagram of the (A) experimental design and (B) analysis pipeline.

### Participants

Data collection took place in “Terra di Unificazione Ewam” center in Florence, Italy. This center is a non-profit, non-sectarian educational institution affiliated with the Foundation for the Preservation of the Mahayana Tradition, specializing in the integration of traditional Tibetan Buddhist psychology and meditative practices within a contemporary secular framework.

All subjects voluntarily participated without any monetary compensation. All participants provided written informed consent to participate in the study. The protocol has been approved by the University of Pisa Ethical Committee N. 26/2023 Prot. 0105602/2023 of 31/07/2023 and conducted in compliance with local regulations, institutional guidelines. All experimental procedures adhered to the principles of Declaration of Helsinki.

Participants were informed about the meditation styles required in the experiment and about the experimental protocol before participation. At the day of the data collection, a brief description was given to each participant at the Centro Terra di Unificazione Ewam, ensuring that they were aware of the types of meditation to be performed and about the overall pipeline of the experiment. Ewam Center was chosen as the place to collect data, since it offers a comfortable, quiet, and meditation-oriented environment, thus limiting discomfort conditions for the participants.

A total of 18 healthy subjects were recruited. Three subjects were removed from the study due to excessive noise and artifacts in the EEG recordings. Thus, the analyzed dataset comprises 15 subjects (mean age ± SD: 49.9 ± 13.6 years; age range: 25–70 years; gender: 9 females, 6 males) with a mean meditation experience of 7.7 ± 7.3 years.

### Experimental Procedure

After sensor placement, the experiment began. Participants were instructed to practice 4-minutes eyes-closed mind-wandering (MW), followed by 15 minutes of each type of eyes-closed meditation: 1. Shamatha (SAM), 2. Vipassana (VIP), and 3. Metta (MET) (see Figure 1 A). We also recorded 2 minutes of resting-state activity with eyes open and eyes closed before and after the meditation sessions; however, these conditions were not included in the present analysis.

MW was performed as a non-meditative control condition (D’Argembeau, 2018; Cahn et al., 2009). During MW, participants were instructed to actively think about autobiographical memories and avoid slipping into a meditative state (**?**). To prevent fatigue, participants were allowed to take short breaks between sessions, resuming the next meditation session. The entire data recording for each participant lasted up to 60 minutes.

### Data collection

EEG signals were acquired using a portable 32-channel electroencephalograph, following the 10–20 international standard montage (EBNeuro BE PLUS PRO FULL, Florence, Italy). Signals were recorded at a sampling rate of 1024 Hz. Participants underwent preliminary scalp scrubbing before wearing the EEG cap. The impedance of all electrodes was measured before the start of the experiment and maintained below 15 kΩ throughout the whole recording. After the experiment, contact resistances of EEG signals were re-checked to collect evidence about any notable differences. A trigger button was attached to the EEG to mark events, allowing data segmentation and labeling across the four experimental conditions.

### Data preprocessing and artifact rejection

Data processing was carried out using EEGLAB (Delorme and Makeig, 2004) open-source software, version 12, running on Matlab R2024a (The MathWorks, Inc.). EEG signals were down-sampled to 256 Hz after applying a low-pass filter with a cut-off frequency of 45 Hz. Subsequently, a high-pass filter at 1 Hz was applied. Bad channels were identified through a two-step semi-automatic procedure. First, channels with a correlation lower than *ρ* = 0.7 compared to their reconstructed versions from neighboring channels were removed (Mullen et al., 2015). This rejection procedure ensured that channel removal was uniform and consistently performed across all the subjects. Then, signals were visually inspected to detect any remaining noisy channel that was not flagged by the correlation criterion (Billeci et al., 2023). Removed channels were reconstructed using spherical spline interpolation (Delorme and Makeig, 2004). After removal of bad channels, the EEG signals were re-referenced to the common average. Independent Component Analysis (ICA) was then applied using the AMICA algorithm (Palmer et al., 2012) to decompose the signals into temporally independent components. The resulting components were automatically classified using ICLabel, and non-brain components with a classification probability greater than 80% were identified as artifacts. These components were further verified through visual inspection of their scalp topographies, power spectral densities, and time courses. Artifact-related components were subsequently removed, and the cleaned EEG signals were reconstructed by back-projecting the remaining components to the sensor space.

MNE Python library was used to epoch data. Specifically, *raw*.*make fixed length epochs* function was exploited to generate non-overlapping 5-second epochs (Shang et al., 2023) from 240 seconds of data for the mind-wandering condition and from 900 seconds of data for the meditation conditions. Consequently, for each subject 48 epochs of mind-wandering and 180 epochs of each of three types of meditation were obtained.

### Feature Extraction & Feature Selection

To characterize the EEG signals, we extracted a comprehensive set of features from three different domains: frequency, time-frequency and time domain features to characterize the EEG signals from the epochs for subsequent classification tasks. From each of the 32 channels, we extracted 29 distinct features, resulting in 928 total features. First, we computed 13 frequency-domain features. To derive these features, we obtained the power spectral density (PSD) using Welch’s method and averaged the PSD values within five frequency bands: *δ* (1–4 Hz), *θ* (4–8 Hz), *α* (8–13 Hz), *β* (14–30 Hz), and *γ* (30–45 Hz). In addition, we extracted spectral entropy, spectral edge frequency, spectral centroid, spectral bandwidth, mean frequency, median frequency, peak frequency, and spectral flatness. Next, we derived 10 time-frequency features using wavelet analysis, “db4” wavelet (Daubechies wavelet) was selected for signal decomposition due to its inherent orthogonality and smoothing properties, which enhance the detection of transient changes in EEG signals (Gowri, 2018). The extracted features consisted of wavelet sub-band energy and wavelet entropy computed across decomposition levels, capturing both the distribution of signal power and the complexity of neural oscillations across frequency bands. Lastly, to quantify the distributional and amplitude-related properties of the signals, we calculated 6 time-domain features including mean, standard deviation, skewness, kurtosis, peak-to-peak amplitude, and root mean square. By integrating these three feature domains, we obtain a richly detailed representation of neural activity, enhancing the robustness and accuracy of the ensuing classification algorithms.

To facilitate effective inter-subject training, features were standardized independently for each subject by subtracting the mean and scaling to unit variance. This normalization reduces inter-individual variability arising from differences in scalp conductivity, electrode impedance, and overall signal amplitude. While such variability may have limited impact in intra-subject classification, it can introduce substantial differences across subjects and negatively affect generalization. In the inter-subject setting, standardization was applied independently to each subject, including the held-out test subject, using statistics computed only from that subject’s data. This approach avoids data leakage while ensuring that subject-specific scaling differences do not bias the classification model.

To improve model efficiency and performance, feature selection was performed using a Random Forest-based approach (Breiman, 2001). Features were ranked according to their contribution to reducing Gini impurity, and a threshold was applied to retain the most informative features. This procedure was implemented within the leave-one-subject-out cross validation (LOSOCV) framework to ensure that feature selection was performed independently in each training fold, thereby avoiding data leakage. By identifying features that were consistently ranked as important across folds, a stable subset of features was obtained for each classification task, reducing dimensionality while preserving the most discriminative information for distinguishing meditative states.

### Classifiers

We evaluated six widely used machine learning classifiers for meditative state classification, including K-Nearest Neighbors (k-NN) (Cover and Hart, 1967), Random Forest (RF) (Breiman, 2001), Support Vector Machines with linear and radial basis function kernels (SVM, SVM-RBF) (Cortes and Vapnik, 1995; Chang and Lin, 2011), Gaussian Naive Bayes (NB) (Bishop, 2006), and a Multilayer Perceptron (MLP) (Rumelhart et al., 1986). All models were implemented using the scikit-learn python library with default hyperparameters unless otherwise specified. For k-NN, the number of neighbors was set to *k* = 5 with Euclidean distance. The RF model used default settings with a fixed random seed `(random state = 42)`. Both SVM variants were implemented using default scikit-learn configurations, with *C* = 1.0 for both models and *γ* = scale for the RBF kernel. The Gaussian NB classifier used default variance smoothing. The MLP consisted of two hidden layers with 100 and 50 neurons, respectively, using ReLU activation and the Adam optimizer.

These classifiers were selected to represent diverse learning paradigms (instance-based, probabilistic, ensemble, margin-based, and neural), enabling a comprehensive evaluation across different decision boundaries. The MLP was designed as a shallow network to serve as a nonlinear baseline while mitigating overfitting given the limited dataset size.

### Intra-subject Training Strategy

To ensure a robust evaluation of classifier performance at the individual level, we adopted an intra-subject training strategy. In intra-subject, training and testing are performed on data from the same subject to evaluate how well meditation states can be detected within subject. Specifically, stratified *k*-fold cross-validation (SKF) with *k* = 10 was applied independently to each subject’s data. In this framework, the data were partitioned into ten mutually exclusive folds while preserving the class distribution within each fold. During each iteration, nine folds were used for training and the remaining fold was held out for testing. This procedure was repeated 10 times, allowing each fold to serve as the test set exactly once. The classification model is therefore evaluated across all folds for each subject, and the resulting accuracies are averaged across subjects to report a reliable estimate of performance.

### Inter-subject Training Strategy

To ensure the classifier generalizes across individuals whose EEG data are not available during training, we adopted an inter-subject (or cross-subject) training strategy. Specifically, we implemented LOSOCV on data from 15 subjects: for each iteration, one of the subject’s epochs were reserved for testing while the remaining 14 subjects’ data were used for training. Consequently, each classification method was tested 15 times on the unseen subject, thereby preventing data leakage between training and testing sets.

### Temporal dynamics of meditation

To examine the temporal stability of classification performance, we conducted a sliding-window analysis. This approach is conceptually similar to single-trial classification methods previously used to track evolving neural states across sessions, for example during therapy-induced recovery (Scaglione et al., 2022). This analysis assesses whether neural discriminability between meditative states changes over time. Starting from meditative versus non-meditative state, a 4-minute mind-wandering segment is slid across 15 minutes of each meditation style separately, using a step size of 50 seconds, yielding 15 time points. Mean classification accuracy is computed at each time point, resulting in accuracy curves over time. For pairwise and multi-class classification, a 4-minute window was selected from each meditation condition and shifted with the same step size to maintain consistency in the temporal analysis. The sliding window is illustrated in Supplementary Material (Figure S1).

We evaluated temporal trends in classification performance by estimating linear regression slopes over time. Specifically, for each classification task, classification accuracy was computed at 15 time points, and a linear regression model was fitted to these values. The resulting slope coefficient *β*_1_ quantifies the direction and magnitude of change in accuracy over time, indicating whether performance increases (*β*_1_ *>* 0) or decreases (*β*_1_ *<* 0) throughout the session. This procedure yields a slope estimate for each subject and classification condition, allowing us to assess temporal trends across all experimental comparisons.

## Statistical Analysis

All statistical analyses were performed in Python using the scipy and statsmodels libraries. Statistical comparison among classifiers performance was conducted using one-way ANOVA with the main factor “Classifier” with levels “NB,” “kNN,” “RF,” “SVM,” “SVM-RBF,” and “MLP”. When a significant main effect was observed, post hoc pairwise comparisons were conducted using Tukey’s honestly significant difference (HSD) test to control for multiple comparisons.

Temporal trends assessed by fitting linear regression models to classification accuracy across time points extracted subject-wise slope coefficients. These coefficients were tested against zero using a one-sample t-test to determine whether changes in performance over time are statistically significant at group level.

For visualization, group-level means are reported along with 95% confidence intervals, computed using the t-distribution. Statistical significance is defined at a threshold of *p <* 0.05.

### Comparison of intra-subject vs inter-subject classification strategy

To compare intra-subject and inter-subject classification strategies, we employ t-distributed stochastic neighbor embedding (t-SNE) to visualize the structure of the learned EEG feature space. t-SNE is a nonlinear dimensionality reduction technique that projects high-dimensional data into a low-dimensional space while preserving local neighborhood relationships, making it particularly suitable for identifying cluster structure in complex neural data (Jeon et al., 2021).

For this analysis, t-SNE embeddings are computed under two conditions. First, intra-subject embeddings are computed using data from a single subject, allowing visualization of subject-specific feature organization. Second, inter-subject embeddings are constructed by pooling data from all subjects into a shared feature space, thereby capturing population-level variability. This dual representation enables a direct qualitative comparison between the two classification schemes.

To further enhance interpretability, a representative subject is highlighted within the pooled embedding. In this setting, the selected subject’s data are visualized alongside data from all other subjects, allowing assessment of if and how individual-specific feature distributions align with or deviate from the global cohort-level structure. All embeddings are computed separately for each classification task, including binary meditative and non-meditative, pairwise, and multi-class conditions.

## RESULTS

We collected EEG data from 15 experienced meditators practicing three distinct meditation styles. We then asked whether EEG signals can reliably distinguish between meditative and non-meditative states (SAM/MW, VIP/MW, MET/MW) and discriminate among different meditation styles (SAM/VIP, SAM/MET, VIP/MET, SAM/VIP/MET, SAM/VIP/MET/MW).

Extracted features were used as input to six classification algorithms: Naive Bayes (NB), k-nearest neighbors (kNN), Random Forest (RF), Support Vector Machine with linear kernel (SVM), Support Vector Machine with radial basis function kernel (SVM-RBF), and multilayer perceptron (MLP). Feature-domain comparisons and feature-selection analyses are provided in the Supplementary Material (Table S1). The Random Forest-selected feature subset achieved the best overall performance and was therefore used in all subsequent analyses.

For inter-subject analysis, we performed 15 repetitions of leave-one-subject-out cross-validation (LOSOCV). For intra-subject analysis, a stratified *k*-fold cross-validation scheme with a 90-10 train-test split was used. We then reported mean ±95% confidence interval accuracies, computed by averaging classification accuracies across repetitions. Chance-level performance was estimated by randomly shuffling class labels and repeating the corresponding cross-validation procedure over 100 iterations.

### Intra-Subject Analysis

Intra-subject classification is conducted to examine within-subject differences across various cognitive states. We first evaluate the recognition between meditative and non-meditative states, as these conditions are expected to be highly distinguishable. Three binary classification tasks were defined: SAM/MW, VIP/MW, and MET/MW. To address class imbalance, equal-length segments are selected, consisting of 4 minutes of meditation and 4 minutes of mind-wandering data. All machine learning models were evaluated and compared to assess whether performance depends on classifier choice. A stratified *k*-fold cross-validation strategy with a 90/10 train-test split is used, and average classification accuracy is computed and reported.

Classification results are presented in Figure 2. Our results showed that meditative states are highly distinguishable from mind-wandering, with accuracies exceeding 90% across classifiers. Among the models, the linear SVM achieves the highest mean accuracy across subjects (SAM/MW: 95.96 ± 1.91%, VIP/MW: 95.78 ± 2.48%, MET/MW: 92.56 ± 3.10%), followed by RF (SAM/MW: 95.15 ± 2.96%, VIP/MW: 95.20 ± 2.48%, MET/MW: 93.75 ± 3.12%).

**Figure 2.**
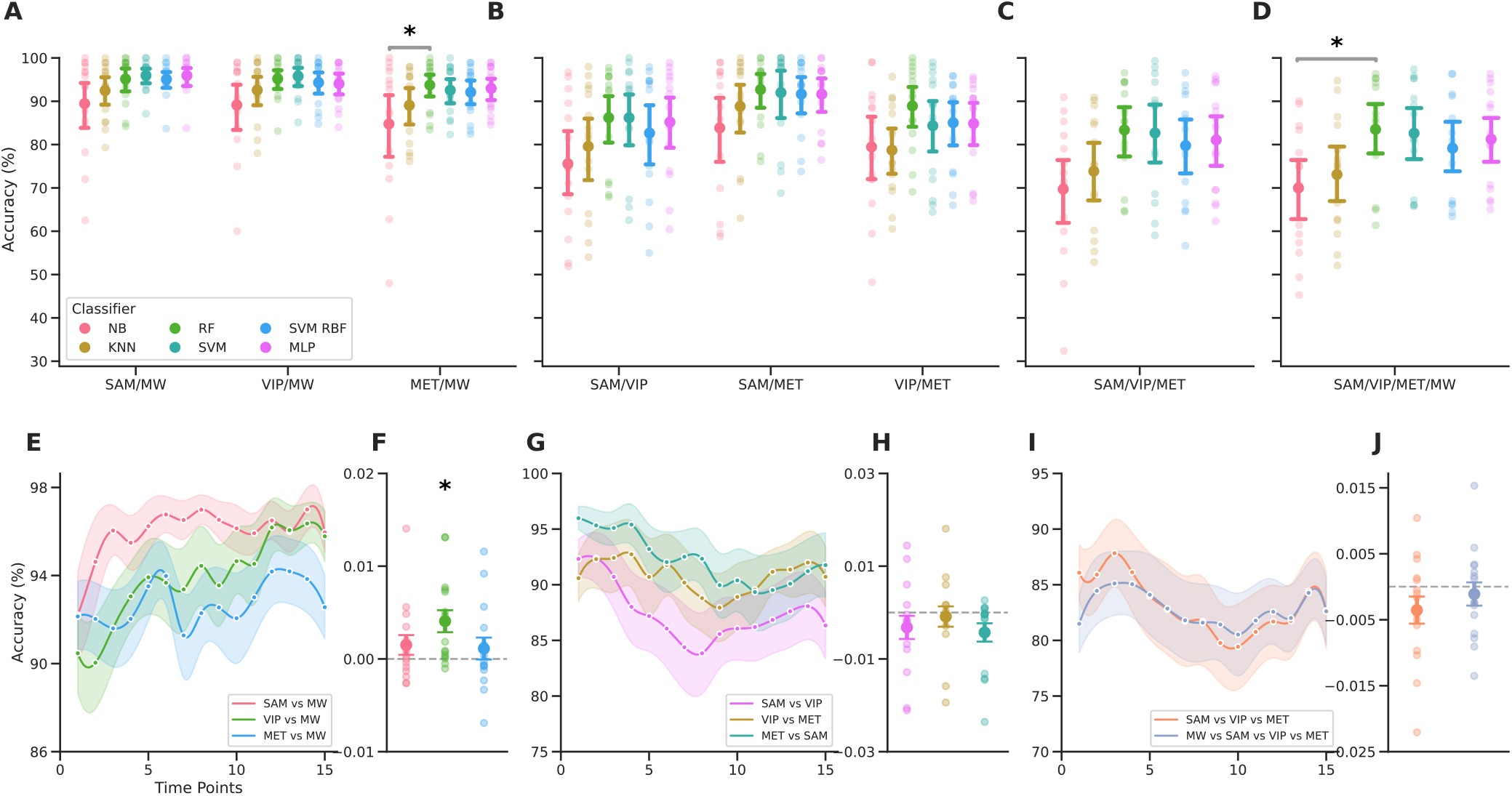
High intra-subject recognition of meditation states and minimal temporal variation. Top row (A–D) shows mean classification accuracies (%) across subjects for multiple tasks and classifiers. (A) Meditation vs. non-meditative state (SAM/MW, VIP/MW, MET/MW) demonstrates consistently high accuracy (*>* 90%), indicating strong within-subject recognition. (B) Pairwise meditation style classification (SAM/VIP, SAM/MET, VIP/MET) also achieves high performance, confirming distinguishable neural features across meditation practices. (C) Three-class (SAM/VIP/MET) and (D) four-class (SAM/VIP/MET/MW) classification show reduced accuracy, reflecting increased task complexity and feature overlap. Error bars denote 95% confidence intervals across subjects, and gray lines indicate statistically significant differences between classifiers. Bottom row (E–J) presents temporal dynamics using sliding-window analysis. (E) Accuracy curves over time for meditation vs. non-meditative state show increasing trends, particularly for VIP vs. MW (shaded regions represent standard error across subjects). (F) Corresponding slope analyses indicate a significant positive temporal trend only for VIP vs. MW (*p* = 0.005, slope = 0.4). (G–J) Pairwise and multi-class meditation style classifications exhibit no significant temporal trends, suggesting stable and non-evolving separation over time.

Building on these findings, we next evaluate the recognition of different meditation styles within subjects. For this purpose, three pairwise classification tasks were performed: SAM/VIP, SAM/MET, and VIP/MET. To ensure consistency across analysis, 4-minute segments are used for each meditation condition. Our results showed that all classifiers demonstrate high discriminability between meditation styles, with RF achieving the highest mean accuracy across subjects (SAM/VIP: 86.21 ± 6.25%, SAM/MET: 92.70 ± 4.33%, VIP/MET: 88.91 ± 5.32%).

To further assess real-world applicability, we evaluate multi-class classification settings. Specifically, three-class (SAM/VIP/MET) and four-class (MW/SAM/VIP/MET) classification tasks were performed. Average accuracies of approximately 80% are achieved across classifiers and subjects. Our results showed that performance decreases with increasing number of classes, reflecting greater overlap in feature space across meditation styles. RF again achieves the highest mean accuracy (SAM/VIP/MET: 83.37 ± 6.09%, SAM/VIP/MET/MW: 83.51 ±6.51%). The complete table of accuracies is presented in Supplementary Material (Table S2).

One-way ANOVA was conducted to evaluate the effect of classifier type on performance across classification tasks. The results indicate that classifier choice exhibits a condition-dependent effect. Significant effects were observed for SAM/MW (*F* = 2.477, *p* = 0.0383), MET/MW (*F* = 2.484, *p* = 0.0378), SAM/VIP/MET (*F* = 2.624, *p* = 0.0296), and SAM/VIP/MET/MW (*F* = 2.951, *p* = 0.0167). In contrast, no significant differences were found for VIP/MW, SAM/VIP, SAM/MET, or VIP/MET. Post-hoc analysis using Tukey’s HSD test showed that the significant effects were primarily driven by differences between the Naïve Bayes (NB) and Random Forest (RF) classifiers. Specifically, NB and RF differed significantly for MET/MW (mean difference = 8.984, *p* = 0.0458) and SAM/VIP/MET/MW (mean difference = 13.5153, *p* = 0.0417).

Temporal stability of classification performance was examined with the sliding-window analysis. The accuracy curves for classification tasks of meditative versus non-meditative state, pairwise meditation style and multi-class meditation state were computed across 15 time points (see Figure 2 E).

Subject-wise regression slopes were analyzed using a one-sample t-test to determine whether classification performance changes significantly over time. A significant positive trend is observed for VIP/MW (*p* = 0.005, *β*_1_ = 0.4), indicating that neural signatures associated with VIP meditation become more stable and more distinguishable over time. In contrast, no significant temporal effects are observed for pairwise or multi-class meditation style classification, suggesting that within-subject separability between meditation styles remains stable throughout the session.

Overall, intra-subject analyses indicate that meditation states are reliably distinguishable with high accuracy. Classification performance is largely robust to classifier choice and temporal variation, suggesting stable and discriminative EEG signatures of meditation within individuals. Based on these findings, we proceed to inter-subject analysis to evaluate the generalizability of learned features and to assess performance on unseen subjects.

### Inter-Subject Analysis

Consistent with the intra-subject analysis, we first evaluate the recognition between meditative and non-meditative states. Classifiers are trained to distinguish meditation from mind-wandering across three meditation styles (SAM, VIP, MET), resulting in three binary classification tasks: SAM/MW, VIP/MW, and MET/MW. We again ensured balanced comparisons by selecting epochs from 4 minutes of meditation and 4 minutes of mind-wandering data.

Classification accuracies are presented in Figure 3. The results demonstrate that meditative states are distinguishable from non-meditative states in unseen subjects, achieving classification accuracies above 70% across classifiers. This indicates the presence of shared neural patterns across participants despite inter-individual variability. Among the evaluated models, SVM-RBF achieves the highest mean accuracy across subjects (SAM/MW: 74.47 ± 8.19%, VIP/MW: 70.72 ± 10.31%, MET/MW: 63.94 ± 7.66%), followed by RF (SAM/MW: 73.88 ± 10.32%, VIP/MW: 69.74 ± 13.94%, MET/MW: 60.33 ± 11.24%).

**Figure 3.**
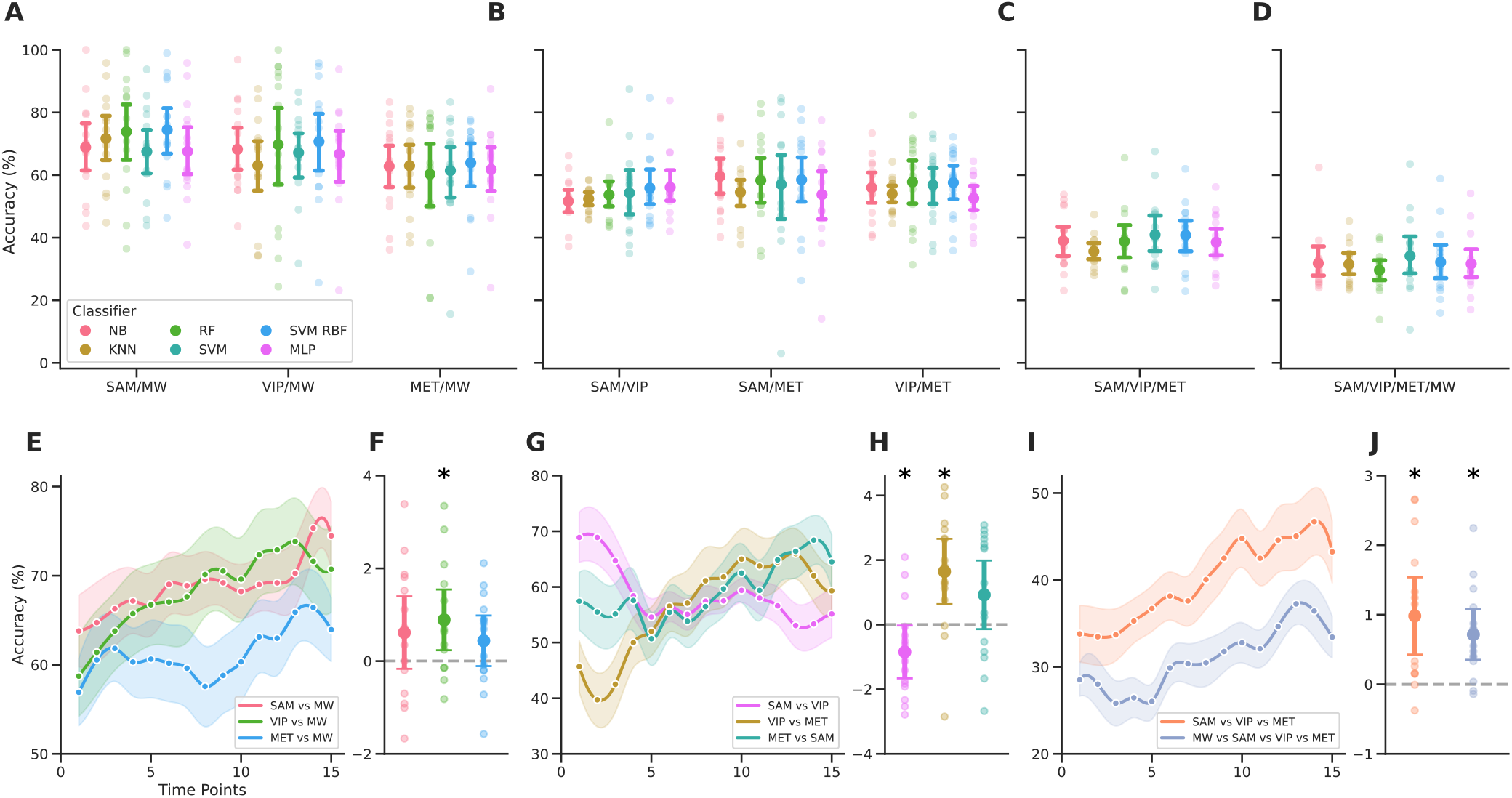
Moderate inter-subject recognition of meditation states with significant temporal variation. Top row (A–D) shows mean classification accuracies (%) across subjects for multiple tasks and classifiers (NB, KNN, RF, SVM, SVM-RBF, MLP). (A) Meditation vs. mind-wandering (SAM/MW, VIP/MW, MET/MW) demonstrates consistently high accuracy, indicating inter-subject recognition. (B) Pairwise meditation style classification (SAM/VIP, SAM/MET, VIP/MET), (C) three-class (SAM/VIP/MET) and (D) four-class (SAM/VIP/MET/MW) classification achieves low but above chance performance, confirming distinguishable neural patterns across meditation practices. Bottom row (E–J) presents temporal dynamics of generalization using sliding-window analysis. (E) Accuracy curves over time for meditation vs. mind-wandering tasks show increasing trends. (F) Corresponding slope analyses indicate a significant positive temporal trend for VIP vs. MW (*p* = 0.011, slope = 0.9). (G–J) Pairwise and multi-class meditation style classifications show significant increasing trends for VIP vs. MET (*p* = 0.004, slope = 1.6) and three (*p* = 0.002, slope = 1.0) and four-class (*p* = 0.001, slope = 0.7), suggesting evolving separability over time.

Building on these findings, we evaluate inter-subject pairwise classification between meditation styles to assess whether distinct neural patterns generalize at the population level. Compared to intra-subject results, classification performance is lower (MLP: SAM/VIP: 56.09±5.64%, SVM-RBF: SAM/MET: 58.50 ±8.58%, RF: VIP/MET: 57.80±7.77%), slightly above chance level (SAM/VIP: 50.17 ±1.03, SAM/MET: 49.99± 1.43, VIP/MET: 49.90 ±1.24), suggesting that although some shared structure exists, meditation styles exhibit substantial inter-individual variability in EEG representations. Multi-class classification performance remains comparatively reduced, performed best with SVM (SAM/VIP/MET: 40.93 ±6.58%, SAM/VIP/MET/MW: 34.15± 6.95%) and above chance level (SAM/VIP/MET: 33.50 ±0.96, SAM/VIP/MET/MW: 24.70 ±1.47). The complete table of accuracies is presented in Supplementary Material (Table S3). No statistically significant differences are observed among classifiers, indicating that classifier choice has minimal influence on inter-subject performance.

To investigate temporal dynamics, we conduct an inter-subject sliding-window analysis using the same configuration as in the intra-subject setting. This analysis evaluates whether generalization performance changes over time during the meditation session. For binary classification, a significant increase in accuracy over time is observed for VIP/MW (*p* = 0.0113, *β*_1_ = 0.9), consistent with intra-subject findings. This suggests that neural features of VIP meditation become more consistent across individuals as the session progresses, improving recognition in unseen subjects.

In contrast, temporal dynamics for meditation style classification are more heterogeneous. A significant decrease in accuracy is observed for SAM/VIP (*p* = 0.044, *β*_1_ = −0.8), whereas significant increases are observed for VIP/MET (*p* = 0.004, *β*_1_ = 1.6) and for multi-class tasks (SAM/VIP/MET: *p* = 0.002, *β*_1_ = 1.0; SAM/VIP/MET/MW: *p* = 0.001, *β*_1_ = 0.7).

These analyses further highlight the differences between intra-subject and inter-subject learning. Although meditation states and styles could still be discriminated in the population level, classification performance remained consistently lower than in the single subject level case. This suggests that meditation-related EEG patterns contain strong subject-specific characteristics, limiting the generalization of classifiers across individuals despite the presence of partially shared neural signatures.

### Feature Space Visualization

We generated both intra-subject and inter-subject embeddings to examine the underlying data structure. In the intra-subject t-SNE visualization, clear and well-separated clusters are observed when the embedding is computed for a single subject subject-05 (see Figure 4), indicating that meditation states are highly separable within individuals, for all eight conditions from meditative and non-meditative state to meditation style comparison and multi-class problem.

**Figure 4.**
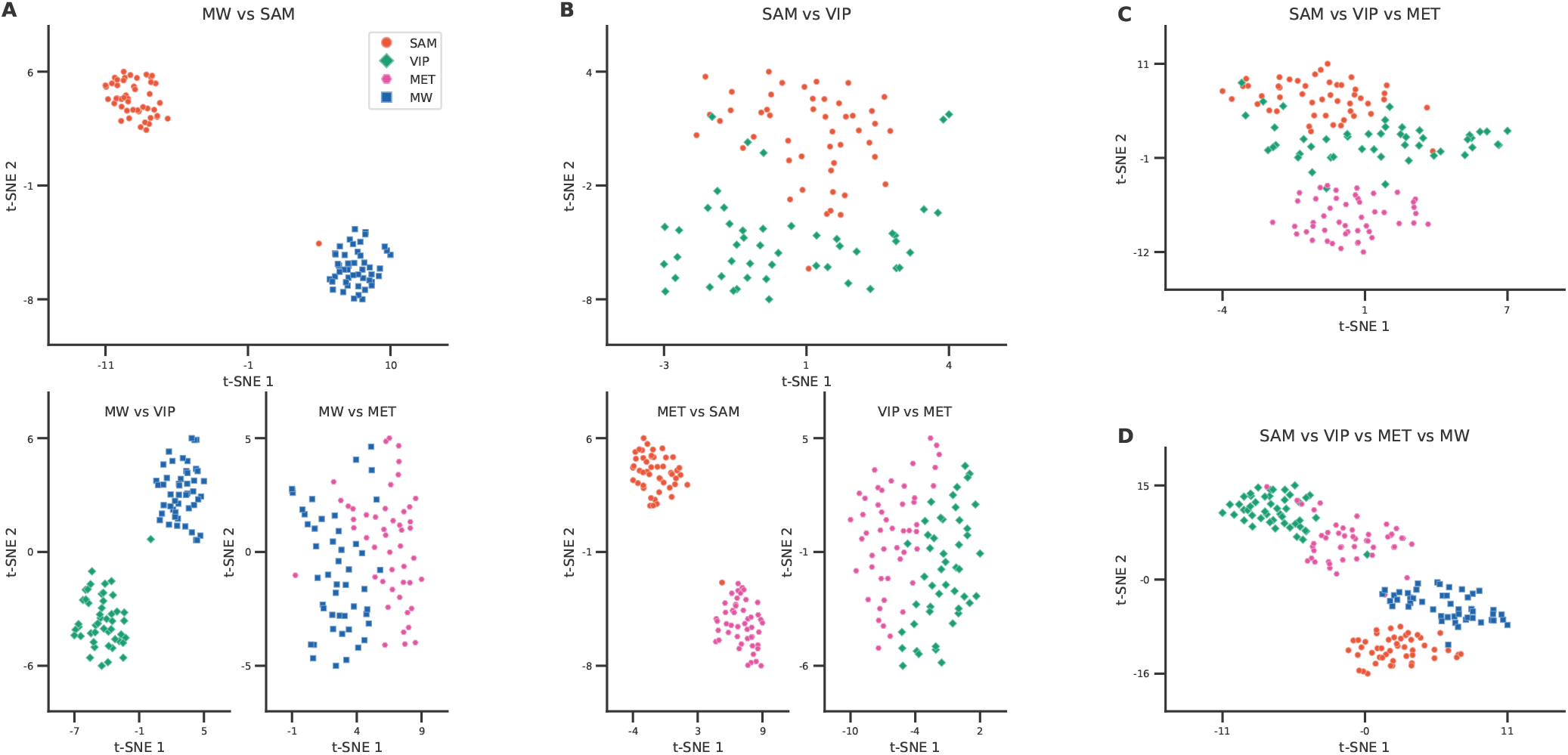
Strong class separability in intra-subject embeddings. t-SNE visualization of EEG features computed for a single subject (subject-05) shows clearly separated clusters across all conditions, including meditation vs. mind-wandering, meditation style comparisons, and multi-class scenarios. This distinct clustering indicates that meditation states are highly separable within individuals, supporting the high classification performance observed in intra-subject models.

In contrast, in the inter-subject embedding, where data from all subjects are pooled into a shared feature space, the clusters become less distinct. While noticeable clusters still appear for SAM vs. MW, VIP vs. MW, and MET vs. MW, the data points show significant overlap for SAM vs. VIP, VIP vs. MET, SAM vs. MET, SAM vs. VIP vs. MET, and SAM vs. VIP vs. MET vs. MW, which explains the lower classification accuracy observed for these tasks (see Figure 5). The relatively clearer separation between meditation and mind-wandering conditions in the inter-subject space explains the moderate performance obtained in these binary classification tasks.

**Figure 5.**
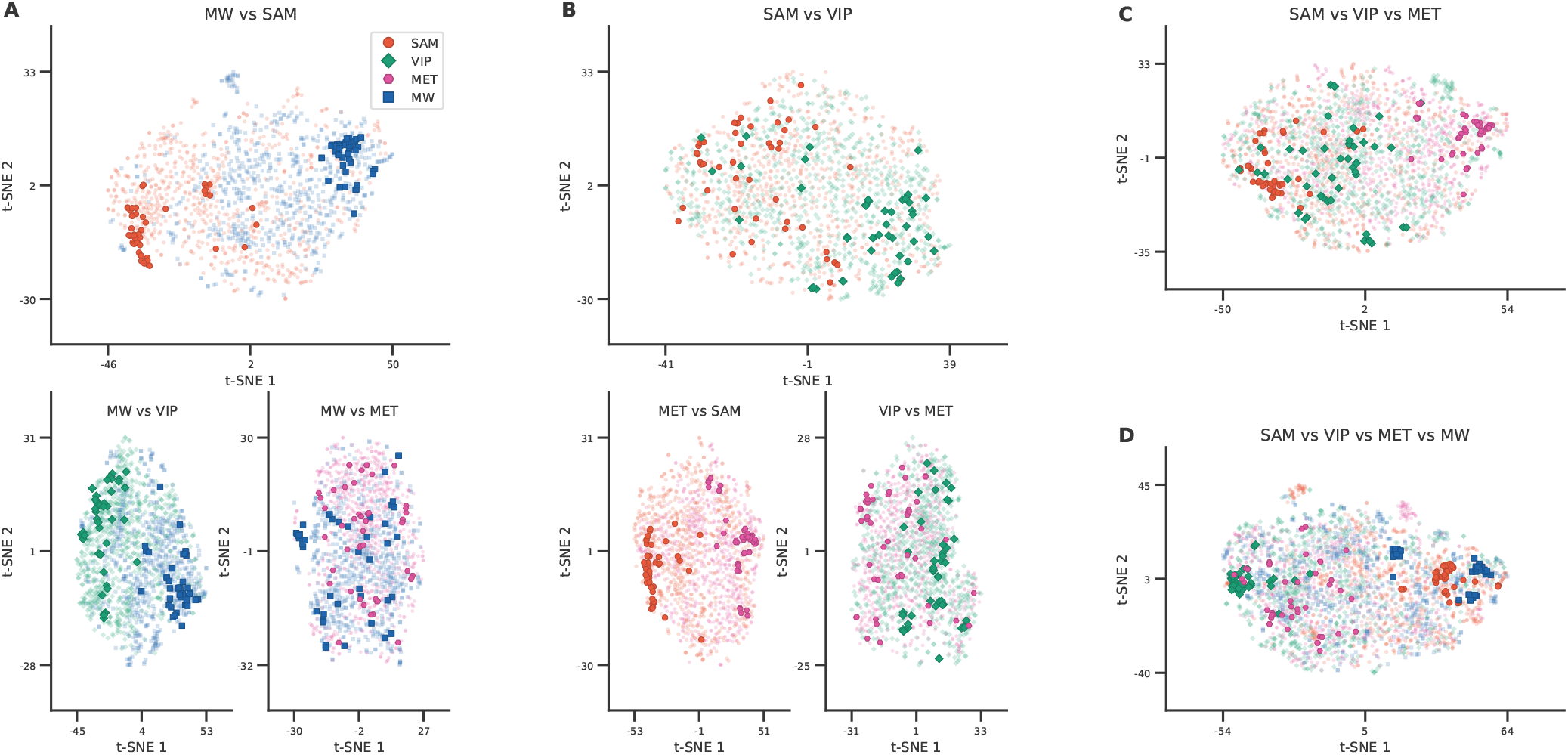
Reduced separability and cluster overlap in inter-subject (pooled) embeddings. t-SNE visualization of pooled space from all subjects reveals that clustering is largely preserved only for meditation vs. mind-wandering conditions (SAM vs. MW, VIP vs. MW, MET vs. MW), while substantial overlap is observed in meditation-style comparisons and multi-class settings. The highlighted points for a single subject (subject-05) retain internally consistent cluster structure, but differ in spatial arrangement from other subjects, leading to mixed feature distributions and reduced classification performance in the shared space.

To further investigate this effect, we highlighted the same individual subject (subject-05) within the pooled embedding space. Consequently, the subject exhibits individually structured clusters corresponding to different meditation states, but these clusters are organized differently for each individual. When data from multiple subjects are combined or pooled together, these subject-specific cluster structures overlap, leading to increased mixing of points in the shared feature space. Methodologically, this suggests that simple feature pooling may not be the optimal way to identify shared neural patterns across subjects; group-level decomposition methods, such as Group ICA, which have been used to extract components common to multiple subjects across brain states (Scaglione et al., 2024), may provide a more principled alternative. For the intra-subject training, the models perform high as the features contain distinct clusters and when the models are trained on multiple subjects for inter-subject training, the models are trained on mixed or confused feature space resulting lower accuracy. This explains why intra-subject models achieve higher performance, whereas inter-subject models experience reduced accuracy due to inter-individual variability in EEG patterns.

## DISCUSSION

The present results demonstrate that EEG-based machine learning can reliably discriminate meditative from non-meditative states at the subject level and with moderate performance at the population level. In contrast, discrimination between different meditation styles achieved high accuracy in the intra-subject setting but remained substantially lower in the inter-subject setting. Feature-space visualization further revealed clear clustering within individuals but increased overlap across subjects, highlighting the influence of inter-individual variability on classification generalization. Additionally, temporal analyses suggest that the discriminability and generalizability of certain meditation states evolve over the course of the meditation session.

### Limitations of the study

This study has several limitations that should be considered when interpreting the results. First, the dataset consists of EEG recordings from 15 Italian meditators with a relatively homogeneous cultural and training background. This may limit the generalizability of the findings and restrict the extent to which inter-subject variability is represented. Future studies should include a larger and more diverse cohort spanning different cultural backgrounds and a wider range of meditation experience, from novice to expert. Such extensions may benefit from multi-center collaborations and larger-scale data collection. The relatively small dataset size also constrains the use of data-intensive approaches such as deep learning, which typically require substantially larger datasets to ensure stable and generalizable training.

Second, the order of meditation styles was fixed due to the inherent structure of the meditation practice. In Tibetan Buddhist traditions, focused attention meditation (i.e., Shamatha) is typically practiced before open monitoring practices such as Vipassana, and Vipassana often precedes loving-kindness (i.e., Metta) meditation (Lutz et al., 2008). Therefore, randomization of condition order was not feasible without altering the naturalistic meditation progression. While this design preserves ecological validity, it introduces a potential confounding factor related to order effects, which may influence neural responses across successive conditions and should be considered when interpreting between-condition differences.

Third, the temporal analysis was conducted on sessions of approximately 15 minutes. The observed time-dependent changes in classification performance may reflect genuine evolution of meditation-related neural states; however, alternative explanations such as within-session adaptation or mental fatigue cannot be fully excluded. As fatigue-related neural changes may overlap with meditation-induced dynamics, future studies should incorporate explicit fatigue monitoring or control conditions to disentangle these effects. Moreover, longer meditation recordings are required to determine whether the observed trends stabilize, continue, or reverse over extended durations.

### Comparison with the state-of-the-art

#### Meditative and non-meditative state classification

Several studies have investigated the classification of meditative versus resting states, typically focusing on a single meditation style, as summarized in Table 1. In contrast, our study evaluates this classification across three distinct meditation styles, providing a more comprehensive and detailed analysis.

**Table 1.** Comparison of classification accuracy across different meditation studies.

| Study | Meditation | Tasks | Method | Inter (%) | Intra (%) | Mix (%) |
| --- | --- | --- | --- | --- | --- | --- |
| Ahani et al. (2014) | Mindfulness | Med/rest | SVM | – | – | 78.00 |
| Han et al. (2020) | Yoga | Med/rest/attention | SVM | – | – | 74.31 |
| Panachakel et al. (2021b) | Raja Yoga | Med/rest | LSTM-α | – | – | 79.10 |
| | | | LSTM-γ | – | – | 94.10 |
| Panachakel et al. (2021a) | Raja Yoga | Med/rest | LDA | 74.00 | – | – |
| Shang et al. (2023) | MBSR | Med/rest | SVM | 64.12 | – | – |
|  |  |  | SCNN | – | 99.65 | – |
| Singh et al. (2025) | Mantra | Med/rest | XGBoost | – | – | 94.73 |
|  |  |  | XGBoost | – | – | 100.00 |
| **Ours** | Shamatha | Med/MW | SVM | **74.47** | 95.96 | – |
|  | Vipassana | Med/MW | SVM | 70.72 | 95.78 | – |
|  | Metta | Med/MW | SVM/RF | 63.94 | 93.75 | – |

Ahani et al. (2014) performed classification of mindfulness meditation in novice participants undergoing short-term Mindfullness-based Stress Reduction (MBSR) training, reporting an accuracy of 78% using an SVM classifier with 10-fold cross-validation. This performance is substantially lower than our intra-subject results, where the SVM classifier achieves 95.78% accuracy for Vipassana meditation. It is important to note that Ahani et al. (2014) employed a mixed-subject cross-validation strategy, where data from all subjects are randomly partitioned into training and testing sets. While this approach can yield optimistic performance estimates, it does not evaluate generalization to unseen individuals and remains dependent on the dataset composition.

This mixed-subject evaluation strategy has been widely adopted in prior studies (Ahani et al., 2014; Han et al., 2020; Singh et al., 2025), limiting direct assessment of model generalizability. In contrast, our study explicitly evaluates both intra-subject and inter-subject scenarios, providing a clearer understanding of subject-specific versus population-level performance.

Panachakel et al. (2021a) performed intra-subject classification using common spatial patterns (CSP) and linear discriminant analysis (LDA) for expert practitioners of Raja yoga (mean experience: 18 years), achieving 97.9% accuracy. This performance is comparable to our intra-subject results (95.96% for Shamatha), despite differences in meditation expertise. Previous work suggests that classification is generally more challenging for novice meditators compared to experts (Han et al., 2020), indicating that our results, obtained from participants with mixed experience (mean experience: 7.7 ±7.3 years), reflect a more challenging and realistic setting.

For inter-subject classification, our SVM-based approach achieves 74.47% accuracy, which is comparable to the 74.0% reported by Panachakel et al. (2021b). In a follow-up study, Panachakel et al. (2021a) report significantly higher accuracy (94.1%) using high gamma features combined with a CSP + LDA + LSTM framework. However, this result is based on a subset of 14 subjects selected from a larger cohort of 54; selection criteria are not explicitly reported, which may introduce bias. Additionally, their study employs eyes-open meditation, which can introduce visual processing and artifacts that confound EEG-based analysis (Cahn and Polich, 2006). In contrast, our study uses eyes-closed conditions, which are generally preferred for obtaining more stable and internally driven neural signals.

Shang et al. (2023) report inter-subject classification accuracy of 68.50% using SVM for novice meditators, while deep learning approaches (ConvNets) perform close to chance level (48.34% and 50.60%). Under comparable experimental settings, including a *5-second* epoch length, our study achieves higher inter-subject accuracy for Shamatha and Vipassana (74.47% and 70.72%, respectively), while Metta (63.94%) remains comparable. For intra-subject classification, Shang et al. (2023) report 98.40% accuracy using shallow ConvNets, which is consistent with the high intra-subject performance observed in our study across all meditation styles.

Singh et al. (2025) employ a mixed-subject classification strategy using XGBoost for mantra meditation, achieving 92.93% accuracy for experienced meditators and 87.59% for novices. However, in this paradigm, participants listen to an external mantra during meditation, which may introduce auditory processing effects. As a result, the classifier may partially capture differences between auditory stimulation and rest, rather than purely internally generated meditation states. In contrast, our study focuses on eyes-closed meditation without external stimuli, ensuring that EEG signals reflect internally generated cognitive processes.

Our results demonstrate competitive or improved performance compared to the state-of-the-art, particularly in inter-subject classification, which remains a challenging and underexplored problem. By leveraging a comprehensive feature set spanning time, frequency, and time-frequency domains across 32 EEG channels, and incorporating feature selection using Random Forest importance, our approach identifies the most informative features for classification. Furthermore, the use of multiple classifiers (NB, kNN, RF, SVM, SVM-RBF, and MLP) ensures robustness of the findings across modeling approaches.

#### Pairwise meditation style classification and multiclass classification

To the best of our knowledge, there is limited prior work that directly performs classification between different meditation styles within the same cohort of subjects. Most existing studies focus on distinguishing meditation from rest or control conditions rather than comparing multiple meditation practices.

Pandey et al. (2022) investigated classification across different meditation groups, including Himalayan Yoga (focused attention), Vipassana (open monitoring), Isha Shoonya (open awareness), and loving-kindness meditation. Their study reports classification accuracies of 84.76% for Himalayan Yoga (Gamma1 band), 90% for Shoonya (Alpha band), and 84.76% for Vipassana (Theta band) when compared against control subjects, highlighting distinct frequency-band characteristics associated with each meditation style. However, their analysis does not directly evaluate pairwise classification between meditation styles within the same experimental setting.

In contrast, our study explicitly evaluates pairwise classification between meditation styles. We observe high intra-subject classification performance across all comparisons, with accuracies of 86.21% for SAM vs. VIP, 92.70% for SAM vs. MET, and 88.91% for VIP vs. MET. In the inter-subject setting, performance decreases (SAM vs. VIP: 56.09%, SAM vs. MET: 58.50%, VIP vs. MET: 57.80%), reflecting increased inter-individual variability. Feature space visualization using t-SNE further supports this observation, showing well-separated clusters within individuals and increased overlap across subjects. Despite this variability, inter-subject performance remains above chance level, indicating the presence of partially shared neural patterns across participants.

Shang et al. (2023) performed multi-class classification using two meditation and two rest conditions, reporting 40.69% accuracy in an inter-subject setting and 79.32% using mixed-subject evaluation. In comparison, although experimental conditions differ, our intra-subject results achieve 83.37% accuracy for three-class (SAM/VIP/MET) and 83.51% for four-class (MW/SAM/VIP/MET) classification. As expected, inter-subject performance decreases but remains above chance level (40.93% for three-class and 34.15% for four-class classification). These findings highlight the increased difficulty of multi-class and cross-subject classification, particularly when distinguishing between multiple meditation styles.

#### Temporal Dynamics of meditation

Previous studies have investigated temporal changes in EEG activity during meditation. Neri et al. (2024) examined EEG dynamics during concentrative and analytical Tibetan meditation using time-varying power spectral density (PSD) analysis. Their results show that EEG features evolve over the course of a session, with more pronounced changes observed in concentrative meditation and among experienced practitioners.

Similarly, Singh et al. (2025) analyzed meditation-related EEG across multiple sessions (before, during, and after a 16-day training period) and reported progressive improvements in classification performance, suggesting that meditation practice influences neural representations over time.

In our sliding-window analysis, we observe temporal effects in classification performance within a single session. Notably, a significant increase in accuracy is observed in inter-subject classification for Vipassana versus mind-wandering over time, based on slope analysis using one-sample t-tests. Similar trends are observed for Vipassana versus Metta and in multi-class classification settings. These results suggest that, as meditation progresses, neural patterns become more consistent across individuals, improving generalizability in inter-subject classification.

## CONCLUSION

This study shows that traditional machine-learning classifiers applied to multi-domain EEG features can reliably distinguish between Shamatha, Vipassana, Metta meditation, and mind-wandering at the within-subject level, with accuracies exceeding 90% for binary meditation-vs-mind-wandering tasks and above 80% for the four-class problem. These results indicate that EEG-based machine learning is a viable tool for personalized assessment of meditative states and could support individualized neurofeedback or progress-tracking applications in which a model is trained on a user’s own data.

In contrast, performance drops substantially in the inter-subject setting, particularly for discrimination between meditation styles, where accuracies approach chance level. Although the obtained results for meditation versus mind-wandering classification in the inter-subject setting are comparable to or slightly better than those reported in prior studies, performance remains insufficient for reliable subject-independent deployment. This pattern, supported by t-SNE visualization of the feature space, suggests that the EEG correlates of meditation are dominated by inter-individual variability and that, with the present feature set and dataset size, it is not yet possible to identify population-level features of meditation suitable for general-purpose, subject-independent meditation-aiding systems. Achieving such generalization will likely require larger and more diverse cohorts, advances in domain adaptation or representation learning, and complementary modalities. For now, the practical pathway toward EEG-based meditation support appears to lie in personalized, within-subject models rather than in universal classifiers.

## AUTHOR CONTRIBUTIONS

SH, FG, PM, AS, and FP conceived the study and designed the protocol. SH, FG, CN, and EC collected the data. SH analyzed the data. SH and AS conceptualized the analyses and generated the figures. SH and AS wrote the first draft of the manuscript. FP provided funding for the study and materials for the study while PM provided the location for the recordings. All authors contributed to manuscript revision, read, and approved the submitted version.

## ACKNOWLEDGMENTS

We acknowledge Mr. Paolo Mori, Centro Terra di Unificazione Ewam Firenze, Istituto Lama Tzong Khapa in Pomaia, and Ente Cassa di Risparmio di Firenze for supporting this study.

## SUPPLEMENTARY DATA

Supplementary tables and figures are provided in the supplementary materials and are referenced throughout the manuscript. These materials contain additional details supporting the analyses and results presented in the main text.

## DATA AVAILABILITY STATEMENT

The raw data supporting the conclusions of this article will be made available by the authors upon reasonable request.

