## Supplementary Material for "Meditation Styles Are Highly Discriminable from EEG at the Subject Level With Limited Generalization Across the Population: A Machine-Learning Study"

### 1 SLIDING WINDOW ANALYSIS

To investigate temporal variations in neural discriminability during meditation, a sliding-window analysis was performed across the full 15-minute meditation session (900 s). A fixed 4-minute analysis window (240 s) was sequentially shifted along the meditation timeline using overlapping temporal segments.

The window positions were distributed across 15 temporal evaluation points (T1–T15), spanning the entire meditation session. The effective step size between consecutive windows was approximately 50 seconds, resulting in substantial temporal overlap between adjacent windows.

At each time point, the window was epoched to 5-second epochs. EEG features were extracted from the corresponding epochs and used for classification analysis. This procedure generated a temporal representation of classification performance, enabling assessment of how neural separability evolved throughout the meditation period (see Figure S1).

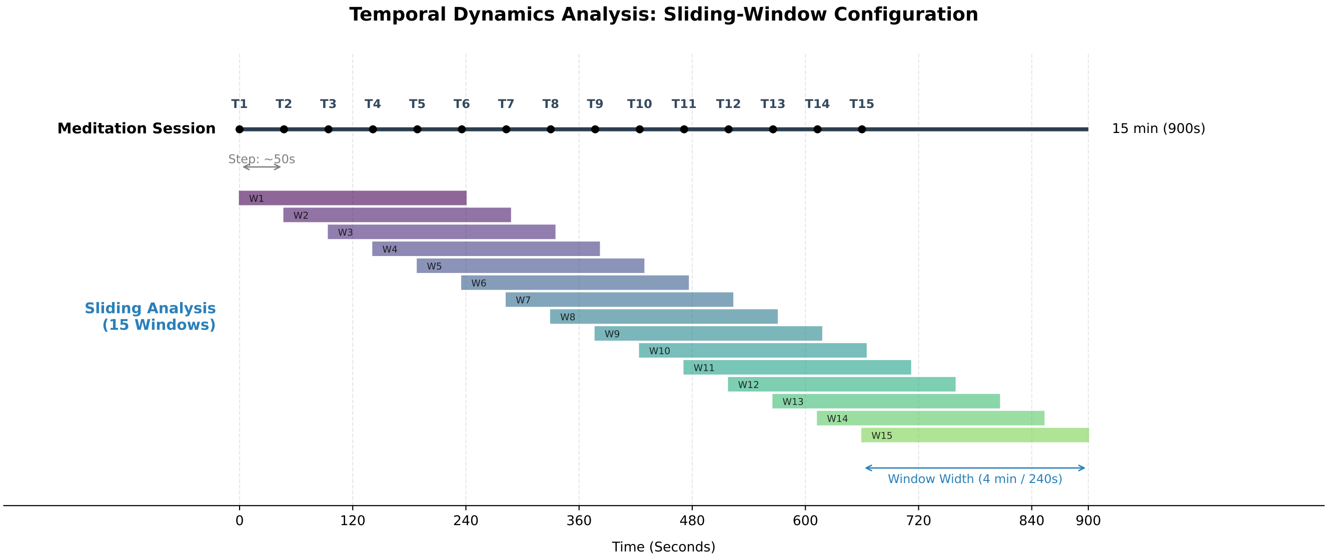


**Figure S1.** Illustration of the sliding-window analysis. A 4-minute window was shifted across the meditation sessions from time points T1 to T15 with an approximate step size of 50 seconds. For meditation versus mind-wandering classification, a fixed 4-minute mind-wandering segment was compared against sliding meditation windows. For pairwise and multi-class meditation-style classification, corresponding sliding windows (W1–W15) were selected from each meditation session.

### 2 EVALUATION OF FEATURE DOMAINS FOR OPTIMAL CLASSIFICATION:

To evaluate the best performing feature set for meditation state recognition, we conducted first classification task (SAM vs. MW) using five distinct feature configurations: Frequency, Time-Frequency, Time, the combined all features (AF) and Selected Features using Random Forest across six classifiers using LOSOCV. The mean accuracies of all 15 models are reported in Table S1. This experiment demonstrated that the Selected Features yielded the best performance across classifiers, with the SVM-RBF classifier achieving the highest accuracy of 74.47%. Therefore, these features were used for all subsequent analysis.

**Table S1.** Classification Accuracy for Different Feature Sets and Classifiers

| Conditions | Features | KNN | RF | NB | SVM | SVM-RBF | MLP |
|---|---|---|---|---|---|---|---|
| SAM vs. MW | Frequency | 60.85 | 63.74 | 60.86 | 52.66 | 63.38 | 58.69 |
| SAM vs. MW | Time-Frequency | 67.61 | 70.82 | 67.68 | 63.89 | 70.77 | 66.64 |
| SAM vs. MW | Time | 55.39 | 62.28 | 60.41 | 59.51 | 64.22 | 55.62 |
| SAM vs. MW | All Features | 62.38 | 71.65 | 66.95 | 58.06 | 69.36 | 67.64 |
| SAM vs. MW | Selected Features | 71.69 | 73.88 | 68.88 | 67.52 | **74.47** | 67.58 |

### 3 INTRA- AND INTER-SUBJECT ANALYSIS

Detailed results of intra-subject analysis are presented in Table S2. Detailed results of inter-subject analysis are presented in Table S3.

**Table S2.** Intra-subject classification performance (Mean ± 95% CI) across classifiers and conditions.

| Condition | Chance | NB | KNN | RF | SVM | SVM-RBF | MLP |
|---|---|---|---|---|---|---|---|
| SAM/MW | 49.61 ± 0.50 | 89.47 ± 6.18 | 92.45 ± 3.62 | 95.15 ± 2.96 | 95.96 ± 1.91 | 95.08 ± 2.18 | 95.92 ± 2.21 |
| VIP/MW | 49.96 ± 0.43 | 89.13 ± 5.94 | 92.58 ± 3.83 | 95.20 ± 2.48 | 95.78 ± 2.48 | 94.32 ± 2.81 | 94.04 ± 2.69 |
| MET/MW | 50.18 ± 0.43 | 84.76 ± 8.30 | 89.05 ± 4.54 | 93.75 ± 3.12 | 92.56 ± 3.10 | 92.09 ± 3.25 | 92.96 ± 2.94 |
| SAM/VIP | 49.68 ± 0.47 | 75.58 ± 8.22 | 79.59 ± 8.04 | 86.21 ± 6.25 | 86.19 ± 6.84 | 82.67 ± 7.74 | 85.21 ± 6.83 |
| SAM/MET | 49.85 ± 0.60 | 83.85 ± 8.19 | 88.83 ± 6.41 | 92.70 ± 4.33 | 91.99 ± 6.00 | 91.65 ± 4.68 | 91.64 ± 4.38 |
| VIP/MET | 49.91 ± 0.63 | 79.46 ± 8.01 | 78.71 ± 5.89 | 88.91 ± 5.32 | 84.35 ± 6.82 | 85.04 ± 5.58 | 84.87 ± 5.54 |
| SAM/VIP/MET | 50.66 ± 0.60 | 69.70 ± 8.79 | 73.85 ± 7.21 | 83.37 ± 6.09 | 82.68 ± 7.42 | 79.77 ± 6.86 | 81.07 ± 6.60 |
| SAM/VIP/MET/MW | 50.16 ± 0.62 | 69.99 ± 8.01 | 73.10 ± 7.33 | 83.51 ± 6.51 | 82.61 ± 6.51 | 79.16 ± 6.48 | 81.25 ± 6.20 |

**Table S3.** Inter-subject classification performance (Mean ± 95% CI) across classifiers and conditions.

| Condition | Chance | NB | KNN | RF | SVM | SVM-RBF | MLP |
|---|---|---|---|---|---|---|---|
| SAM/MW | 50.06 ± 2.46 | 68.88 ± 8.35 | 71.69 ± 7.94 | 73.88 ± 10.32 | 67.52 ± 7.90 | 74.47 ± 8.19 | 67.58 ± 8.54 |
| VIP/MW | 49.39 ± 2.59 | 68.24 ± 7.86 | 63.06 ± 9.20 | 69.74 ± 13.94 | 67.16 ± 7.57 | 70.72 ± 10.31 | 66.74 ± 8.72 |
| MET/MW | 49.59 ± 2.22 | 62.81 ± 7.71 | 63.00 ± 7.85 | 60.33 ± 11.24 | 61.44 ± 9.06 | 63.94 ± 7.66 | 61.80 ± 7.93 |
| SAM/VIP | 50.17 ± 1.03 | 51.64 ± 4.13 | 52.40 ± 2.34 | 53.68 ± 4.63 | 54.31 ± 7.68 | 55.87 ± 6.31 | 56.09 ± 5.64 |
| SAM/MET | 49.99 ± 1.43 | 59.59 ± 6.56 | 54.54 ± 4.65 | 58.30 ± 7.93 | 57.01 ± 11.40 | 58.50 ± 8.58 | 53.76 ± 8.68 |
| VIP/MET | 49.90 ± 1.24 | 55.98 ± 5.25 | 54.01 ± 3.08 | 57.80 ± 7.77 | 56.86 ± 6.33 | 57.59 ± 6.20 | 52.61 ± 4.33 |
| SAM/VIP/MET | 33.50 ± 0.96 | 39.01 ± 5.29 | 35.61 ± 2.83 | 38.82 ± 5.88 | 40.93 ± 6.58 | 40.81 ± 5.62 | 38.56 ± 4.91 |
| SAM/VIP/MET/MW | 24.70 ± 1.47 | 31.82 ± 5.19 | 31.52 ± 3.56 | 29.63 ± 3.63 | 34.15 ± 6.95 | 32.20 ± 6.14 | 31.65 ± 5.27 |
